# Accounting for demography in the assessment of wild animal welfare

**DOI:** 10.1101/819565

**Authors:** Luke B.B. Hecht

**Affiliations:** Department of Biosciences, Durham University, Durham, United Kingdom

## Abstract

Welfare is experienced by individual animals, but the quantity and average quality of welfare an individual is likely to experience in their lifetime is bounded by population demography; namely, age-specific survivorship and the ecological forces that shape it. In many species, a minority of the individuals who are born survive to adulthood, meaning that the lives of those we observe in nature are often unrepresentative of the typical individual born into their population. Since only living animals are capable of experiencing welfare, lifespan is effectively an upper bound on the amount of affectively positive or negative experience an animal can accrue. Life history strategies that increase the probability of a long life are therefore more permissive of good welfare; but even holding life expectancy constant, specific patterns of age-specific mortality may enable a larger proportion of individuals to live through periods characterized by above-average welfare. I formalize this association between demography and welfare through the concept of welfare expectancy, which is applied to published demographic models for >80 species to illustrate the diversity of age-specific mortality patterns and entertain hypotheses about the relationship between demography and welfare.

## 1. Introduction

The experiences of wild animals are extraordinarily diverse. Individuals of different species occupy different habitats, consume different resources, and engage in different behaviors. Even within species, animals’ fortunes differ based on their relative fitness or due to chance events, leading to differential survival or mating success. While life history strategies evolve to maximize inclusive fitness, it is crucial to recognize that fitness and welfare are not the same (Beausoleil et al., 2018). For example, a strategy which maximizes mean fitness may do so while increasing the variance in outcomes among siblings, leading to reproductive success in adulthood for a few, but short lives for most (Pettorelli and Durant, 2007). Even for a successful individual, high evolutionary fitness need not imply good welfare, as sexual competition forces trade-offs between reproduction and survival or somatic maintenance (Johnston et al., 2013).

A key objective of the nascent fields of conservation welfare (Beausoleil et al., 2018) and welfare biology (Ng, 1995) is to evaluate the quality of lives lived by wild animals in order to identify causes of poor welfare, as well as safe and tractable interventions to improve welfare. Empirical evaluations of wild animal welfare, such as those based on stress hormone levels and other veterinary techniques, have been carried out, though few are comparable between contexts (Schwarzenberger, 2007). One of the most promising proposals to date is the use of differential rates of biological aging as an indicator of lifetime cumulative welfare under different conditions (Bateson and Poirier, 2019; Poirier et al., 2019). For example, social stress related to brood size and social rank has lifelong fitness consequences in birds that appear to be mediated by telomere attrition, a prominent biomarker of biological age (Boonekamp et al., 2014; Nettle et al., 2015). More generalized application of these methods will require hypotheses to test and a framework for prioritizing which populations to evaluate and which groups of individuals within them to potentially aid.

A promising starting point is to reason from demographic patterns such as the distribution of individual lifespans. This approach is compatible with a wide range of assumptions about the causes and levels of wild animal welfare, since only living animals are capable of experiencing welfare. Moreover, the quality of life experienced by a typical week-old animal is likely different from that of a year-old animal due to changing levels of vulnerability to disease and predation, competition with conspecifics, self-sufficiency, and senescence.

Demographic patterns are observed at the population level but experienced by individuals. Population models can provide examples of potential demographies, and although no firm conclusions about welfare can be drawn from interspecific comparisons given our uncertainty about the preferences and experiences of most animals, we can use their diverse population dynamics to probe the implications of different hypotheses for how welfare varies with age.

Here, I set out a framework for incorporating demography in the evaluation of wild animal welfare based on the principle of expected value and formalize previously expressed intuitions about the relationship between life history and welfare. I also illustrate this by application to published matrix population models for 160 populations of >80 species and formulate working hypotheses about welfare to be tested by future field studies.

## 2. Methods

### 2.1 Matrix population models

Matrix population models (MPMs) use matrix algebra to represent transitions between life stages in a population (Caswell, 2001). They are widely used to infer populations’ instantaneous rates of growth, as well as for estimating vital rates such as survival and fecundity, and conducting population viability analyses (Heppell et al., 2000). MPMs may be structured by age, stage, or another trait, such as size. For the analyses described here, I used exclusively age-based models to avoid any ambiguity around stage durations.

It is possible to calculate the expected distribution of lifespans among a cohort from the age-specific mortality rates given by an MPM. Life expectancy is then calculated as the sum of the probability of each possible lifespan multiplied by its length (Caswell, 2009). Annual survival probabilities were assumed to be the product of equal daily survival probabilities, so individuals dying during a given year were assumed to have lived through half of that year.

### 2.2 Welfare expectancy

Life expectancy from birth (*e*_*0*_) represents the expected value of lifespan, with the “value” of each possible lifespan being equal to its length, each additional year of life being weighed the same. Calculations of generation time – the expected age of mothers – follow a similar formula but allow ages to differ in value as age-specific fecundity varies. Welfare may similarly vary with age, as juveniles, sub-adults, reproductive adults and senescent animals face different levels and forms of disease, competition, predation and environmental hardship. This potential for variation calls for a distinct concept of welfare expectancy.

Welfare expectancy from birth (*W*_*0*_) is calculated by summing the age-specific welfare values experienced over the ages encompassed by each possible lifespan, and then taking the mean weighted by the probability of each lifespan: 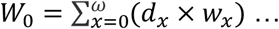 where *d*_*x*_ = probability from birth of dying at age *x*; *w*_*x*_ = net total welfare experienced during a lifespan of *x* years; ω = maximum lifespan. For example, the expected value of a 5-year life would be equal to the total amount of welfare experienced between ages 0 and 5, multiplied by the probability of a 5-year lifespan. Repeating this operation for each possible lifespan and taking the sum would yield the welfare expectancy for a typical individual born into the population in question.

A relative welfare expectancy (RWE) index can also be calculated using values of *w*_*x*_ normalized around one 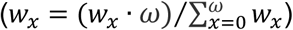 to calculate welfare expectancy (*W*_*0,R*_), and then dividing welfare expectancy by life expectancy: RWE = *W*_*0,R*_/*e*_*0*_. This index expresses the variability of lifespan in relation to periods of high or low welfare. An RWE > 1 implies that most individuals will survive to experience periods of life characterized by above-average welfare, while for RWE < 1, a population’s survivorship patterns mean that most individuals will only experience below-average periods of life. For RWE > 1, welfare can be said to be ‘outperforming’ life expectancy, as the average instantaneous welfare value during the lifetime of a typical individual would be greater than for an individual who lived out their theoretical maximum lifespan. In general, either high early-life survivorship/welfare (many individuals experiencing the best years) or extremely low late-life survivorship/welfare (few individuals experiencing the worst years) can yield a high RWE index. As life expectancy approaches the maximum lifespan of a species, RWE will tend towards 1 because the average welfare an individual experiences in their lifetime is increasingly representative of the welfare distribution over their species’ maximum lifespan.

### 2.3 Welfare elasticity analysis

An elasticity analysis was also applied to each population to see whether they differed in the age at which a proportional reduction in mortality rate would have the greatest impact on individual’s lifetime welfare expectancy. The elasticity of welfare expectancy to each age’s survival rate was scored as the product of 1) the survivorship up to that age (*l*_*x*_), 2) the mortality rate at that age (*m*_*x*_), and 3) the remaining welfare expectancy conditional on surviving that age (*W*_*x*_). The age with the highest elasticity score was considered the welfare ‘bottleneck’ age for individuals of that population.

### 2.4 Illustrating the welfare expectancy approach

To provide an initial illustration of the approach described here using as detailed and explicit a case as possible, a Leslie matrix was generated from age-specific rates of survival and welfare (life satisfaction) among the human population of the United Kingdom, using published statistics from the UK Office for National Statistics (ONS, 2016; 2019). This population was subjected to the welfare expectancy analyses described above. It is so far unique in having empirically determined age-specific welfare values, as well as vital rates calculated from known fates of thousands of individuals, permitting the clearest possible illustration of the welfare expectancy approach.

The UK human lifespan distribution begins with a modest spike representing infant mortality. It then abruptly falls after the first year, rising again gradually from throughout senescent life before spiking at 88 (Figure 1). The life expectancy at birth was approximately 80 years. Welfare (life satisfaction) is bimodal, with peaks in the early-twenties (beginning of independent life) and mid-sixties (beginning of retirement) and troughs in the mid-forties and old age. The population’s RWE index was 1.00, as the vast majority of individuals lived to old age and experienced periods of high and low welfare in roughly equal measure (Figure 1, left). The age at which welfare expectancy was most elastic to a marginal reduction in mortality was during year 1, combating low but non-trivial infant mortality (Figure 1, right). This is to be expected given that all individuals are alive and able to benefit from interventions at this age, and individuals surviving infanthood may expect a long and happy life. Notably, age 80 has only slightly lower elasticity. This is because, although welfare expectancy from age 80 onward is much lower than welfare expectancy from birth, the population’s extremely high survival rates up to old age mean that ∼60% of individuals survive to benefit from interventions at age 80. Moreover, because the age-specific mortality rate is much higher than during infanthood, any intervention may have a proportionally greater effect.

**Figure 1:**
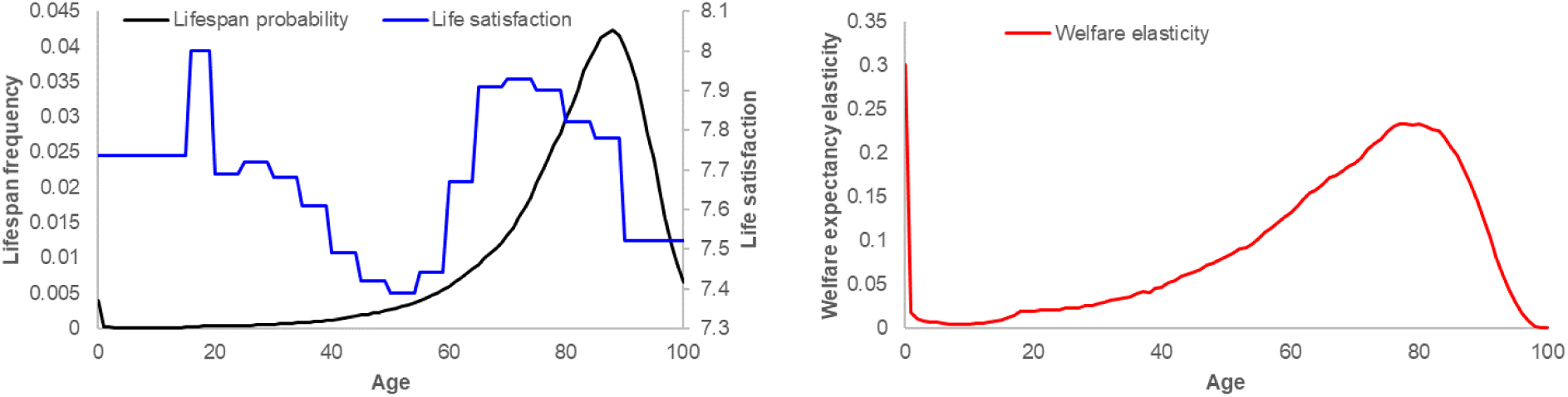
(Left) The lifespan distribution of UK humans plotted against age-specific welfare (life satisfaction). Most individuals die old enough to experience both highs and lows of welfare coinciding with important life transitions. (Right) Elasticity of welfare expectancy to marginal reduction in age-specific mortality rate.

### 2.5 Modelling age-specific welfare

The distribution of welfare with respect to age is a crucial determinant of how changes in demographic vital rates affect individual welfare expectancy, but there is yet virtually no direct evidence on the age-specific welfare of wild animals. However, to explore the implications of varying age-specific welfare, I assumed that welfare at a given age was proportional to the probability of surviving that year of life. It must be stressed that this is a working hypothesis, adopted for the purpose of illustrating the effects of age-dependent welfare under various real-life demographies. The assumption remains to be tested, but its rationale, implications and alternatives will be discussed later. Welfare expectancy specifically calculated under this assumption will be denoted as *W*_*0,S*_.

### 2.6 Data obtention

Published MPMs were obtained from the COMADRE database, which serves as a curated repository for matrix population models (Salguero-Gomez et al., 2016). A subset of 152 population matrices, representing 88 species, were selected according to the following criteria, in the form of variables defined in the COMADRE documentation: MatrixComposite == Mean & MatrixTreatment == ‘Unmanipulated’ & MatrixCaptivity == ‘W’ & MatrixSplit == Divided & ProjectionInterval == 1 & MatrixCriteriaOntogeny == ‘No’ & MatrixCriteriaSize == ‘No’ & MatrixCriteriaAge == ‘Yes’. Only the survival matrices ($matU) were used. From this subset, matrices were discarded if they had missing data (“NA” values), stage-specific transition probabilities summing to >1 or to 0 at non-terminal stages or were duplicates. All MPMs were annual Leslie or Leslie+ matrices (Carslake et al., 2009). Original credit for these matrices goes to their respective authors, as attributed in the COMADRE database.

Four major taxonomic classes were represented among the population matrices drawn from COMADRE: Actinopterygii (ray-finned fishes), Aves (birds), Mammalia (mammals), and Reptilia (reptiles). These were represented by 16, 54, 72, and 10 populations, respectively. Maximum lifespans for each species was obtained from the AnAge database (De Magalhães et al., 2005), if available, or else imputed as the average of represented congeners or family relatives. In the case of Leslie matrices, the maximum lifespan was determined by the dimension of the matrix itself. Statistics for each of these matrices can be found in appendix Table A1.

## 3. Results

### 3.1 Life expectancy

The mean life expectancy across the wild animal population models obtained from COMADRE was calculated at 4.39 years, or a median of 3.14 years. Approximately 16% of populations had life expectancies of <1 year, and 74% had life expectancies of <5 years. As a proportion of maximum lifespan, the average life expectancy was 16%, with only 5% of populations having life expectancies >33% of their maximum. Mammal populations had the highest average life expectancy (6.8 years), followed by birds (2.8 years) and reptiles (2.0 years). The ray-finned fish had the lowest average life expectancy, at 0.6 years (Figure 2).

**Figure 2:**
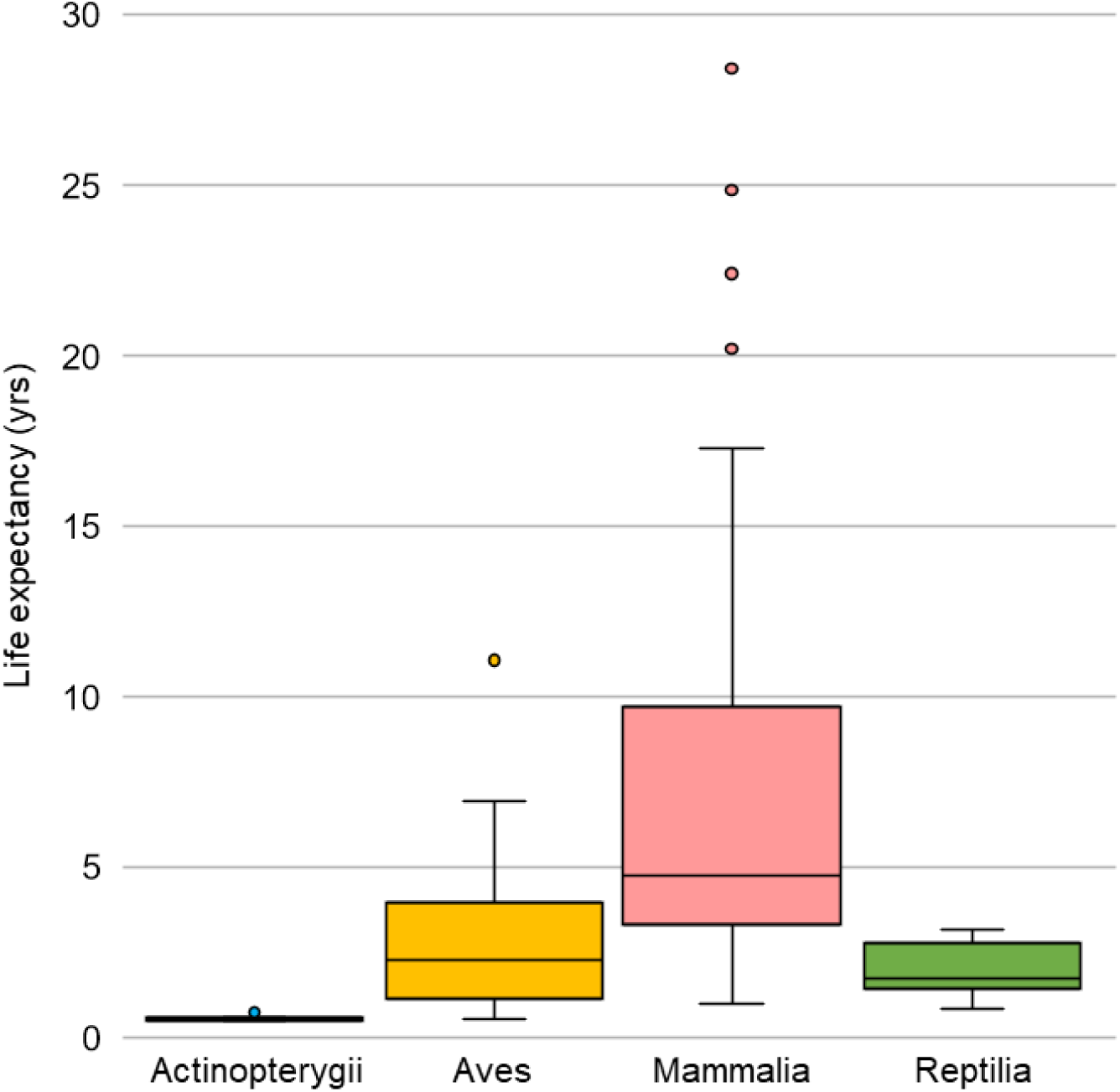
Box and whiskers plot of life expectancy by taxonomic class. Populations were used as data points.

### 3.2 Welfare expectancy

The mean RWE index was 0.87, median 0.97. Only 10% of populations had relative welfare expectancies of >1.1, while 33% scored <0.9. Mammal and bird populations had RWE values typically near 1 (Figure 3). Mammalia had a tighter distribution around 1, consistent with the longer life expectancies of mammalian populations, but with positive and negative outliers. Actinopterygii had by far the lowest mean RWE (0.20). Reptilian RWE values were intermediate, though all below 1 except for an extreme positive outlier (RWE=2.38) based on data from a population of painted turtles (*Chrysemys picta*) at E.S. George Reserve (Tinkle et al., 1981).

**Figure 3:**
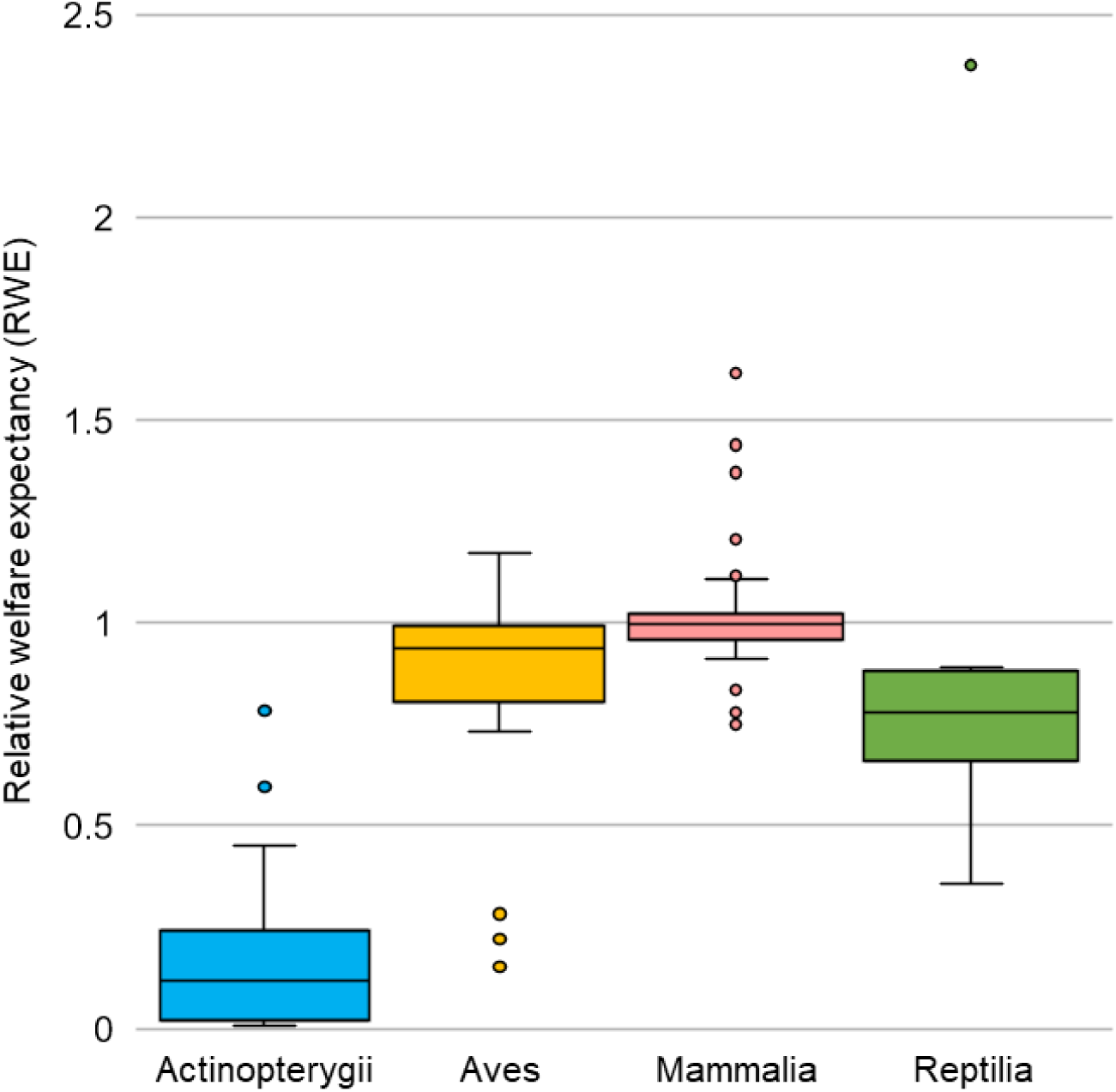
Box and whiskers plot of RWE by taxonomic class, using populations as data points. The high-RWE outlier among Reptilia is a population of painted turtles (*Chrysemys picta*).

The average annual survival distributions of all populations with low (first quartile) and high (third quartile) RWE were plotted and found to cover distinct value ranges only during early life. High-RWE populations sustained a relatively high survival rate from birth onwards. Meanwhile, low-RWE populations had extremely low first-year survival rates, yet many attained higher survival rates similar to those of high-RWE populations by age 6 (Figure 4).

**Figure 4:**
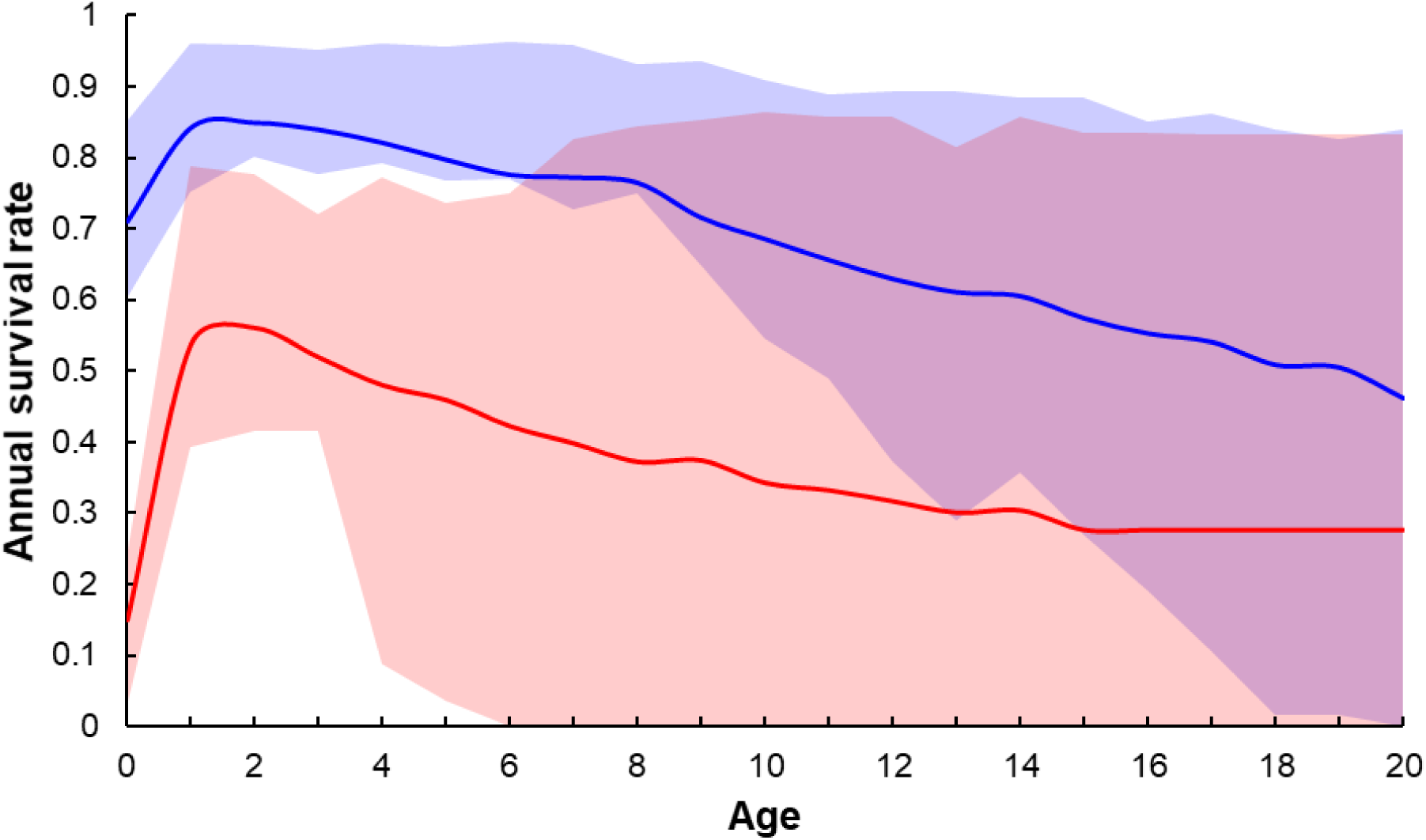
Mean age-specific annual survival rates for high-RWE populations (top 25%; blue line) and low-RWE populations (bottom 25%; red line). Lighter blue and red bands cover the interquartile ranges about each mean line. The purple area depicts overlapping age-specific survival distributions of the high- and low-RWE groups.

### 3.3 Age-specific elasticity of welfare expectancy

The elasticity analysis identified only nine populations for which an infinitesimal reduction in mortality rate after age 0 would lead to a greater increase in welfare expectancy than an equivalent reduction in first-year mortality. For five of these populations, the age of highest welfare elasticity was year 1 or 2, enabled by high survivorship over the preceding period followed by a drop (the ‘bottleneck’). The other four bottlenecked populations belong to the same species, *Capra ibex*, and show a distinct lifespan distribution that leads to peak welfare elasticity around age 7 or 12. In both cases, the elasticity of welfare expectancy to an age-specific reduction in mortality parallels the lifespan distribution (Figure 5).

**Figure 5:**
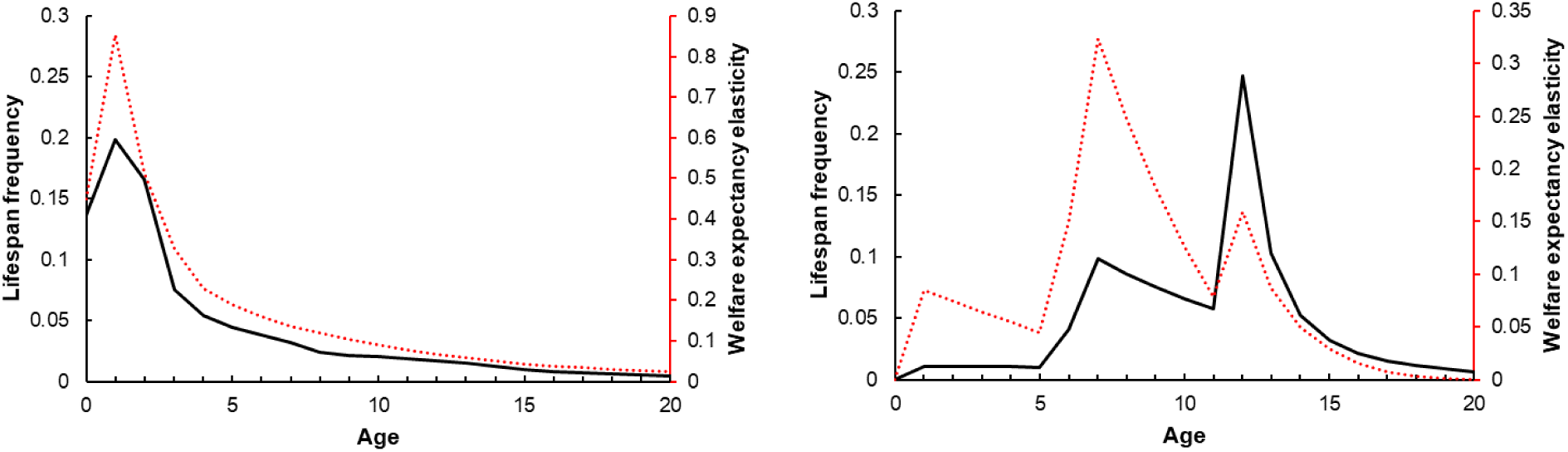
The lifespan distribution (solid black line) and corresponding age-specific welfare elasticity (dotted red line) of five ‘bottlenecked’ populations with low age 0-1 survival but higher age 1-2 and 2-3 survival (left) and four *Capra ibex* populations where welfare elasticity peaked around age 7 or 12 (right).

## 4. Discussion

### 4.1 Life expectancy

Most of the individuals observable at any given time in many wild animal populations are the lucky ones who have survived the challenges characteristic of early life. Among the populations considered here, based on published demographic models, the typical life expectancy is merely 14% of the theoretical lifespan. While this is the median across populations, given that predominantly shorter-lived taxa, such as the ray-finned fish, may produce far more offspring per generation than longer-lived ones, the average life expectancy across individuals is likely to be much smaller. The criterion of annual periodicity used for selecting population matrices from COMADRE could further bias life expectancy upward, since an annual time-step would provide poor resolution when studying a very short-lived animal. This is particularly relevant for considering the lifespans of juveniles, which may encompass a fraction of one year.

Not all newborns of a given population will have the same individual life expectancies, after the predictive power of parental phenotypes and circumstances of birth are taken into account. Parental age, maternal body mass, clutch size, and relative timing of birth have often been found to predict lifespan (e.g. Reid et al., 2010; Einum and Fleming, 2000; Tamada and Iwata, 2005; Ronget et al., 2018). Field research developing such predictors of individual differences could help define life expectancy more precisely for subsets of a population, helping to target interventions on the most vulnerable animals.

### 4.2 Age-specific welfare

Variation in lifespan also magnifies the relevance of differences in average quality of life with respect to age among a given population. In a comparison of two populations with the same life expectancy and theoretical lifespan, the one in which the largest proportion of individuals survive to experience the most pleasant years of life available to them will have a greater potential for net-positive welfare.

In the present analyses, I assumed that the welfare experienced at a particular age was proportional to the probability of surviving that year of life. This is a plausible working hypothesis since the same factors that lead to mortality (e.g. disease, vulnerability to predators, competition for food) have been shown to lead to chronic stress and poor physical condition (Clinchy et al., 2013; Bateson et al., 2015). Assuming this model of age-specific welfare, and equal life expectancies, populations with a) very low mortality in early life followed by high mortality later in life would achieve higher welfare than populations with b) a constant rate of mortality, and these would in turn achieve higher welfare than c) populations with high early-life mortality but high adult survivorship. These scenarios roughly correspond to the survivorship curve typology of Demetrius (1978).

A number of alternative hypotheses might also describe the relationship between welfare and age. For example, welfare might peak around the same age as peak reproduction. This could occur due to hormonal factors, or simply because natural selection tends to optimize fitness around reproduction, and body condition is likely related to welfare; though this might be perturbed by intense juvenile competition or the need to provide protection for offspring, which could drive peak physical fitness earlier or later than peak fecundity. On the other hand, reproductive age might bring on poor welfare, especially in species with intense sexual competition (e.g. Clinton and Le Boeuf, 1993). Either of these reproductive age-centric hypotheses would likely still predict a correlation between survival and welfare, given the interaction of age-specific mortality and reproductive timing in directing the evolution of life history strategies (Charlesworth, 1980).

It is also conceivable that the determinants of welfare are so complex that welfare varies irregularly over a lifetime, or average welfare might even be invariant with age in some animals. If welfare is invariant with age, welfare expectancy will scale linearly with life expectancy. However, it seems highly likely that welfare would shift in some direction concurrent with major life history transitions, like the maturation of a tadpole or caterpillar, or sexual maturation in most species, especially when this is accompanied by changes in environment, such as with the ejection of young male hyenas or female meerkats from their natal groups (Maag et al., 2019).

Previous reviews have recognized the need to integrate welfare experienced over the lifetime of domestic animals (e.g. FAWC, 2009; Pickard, 2013). The concept of welfare expectancy developed here applies this to wild animal populations, using the principle of expected value to account for their inherent variability. Recently, Bateson and Poirier (2019) proposed that the ratio between biological and chronological age could be used as a proxy for lifetime welfare. The premise of this approach is that somatic damage and repair, which determine biological age, often result from physiological processes that are associated with affective states, such as stress or happiness. Indeed, adverse conditions such as sibling competition have been shown to lead to accelerated biological aging limited to the study period, especially when the individual is a weaker competitor (Gott et al., 2018). Surveying population-level variation or tracking individual longitudinal variation in the biological-to-chronological age ratio, through measurements such as telomere length, could be a cost-effective way to estimate relative age-specific welfare within wild populations. In the Anthropocene, a large proportion of wild animal stress may be caused by human activity, and so biomarkers such as these could provide evidence of habitat quality from the perspective of the animals themselves and serve as additional holistic evidence to present policymakers (Wikelski and Cooke, 2006).

### 4.3 Welfare expectancy

Since only living animals are capable of experiencing any level of welfare, life expectancy has profound implications for the net welfare of a population. I have defined welfare expectancy from the perspective of an individual being born into a population and facing an uncertain lifespan. Welfare expectancy revolves around age-specific variation in welfare and the implication that some lifespans will encompass a greater quality and quantity of welfare than others. Many animals die as juveniles, only experiencing the level of welfare associated with that stage of life as a member of their species; others survive to adulthood but fail to reproduce, while others live long, iteroparous lives.

The potential for age-specific variation in average welfare suggests that welfare expectancy may ‘outperform’ life expectancy in populations where welfare is highest in early life, which most individuals will live to experience. Conversely, in populations where juvenile welfare is lower than adult welfare, welfare expectancy may ‘underperform’ life expectancy because most individuals never see their best years. This notion drives the concept of relative welfare expectancy (RWE). Assuming the correlation between age-specific survival and welfare argued above, welfare expectancy in one third of the populations considered here underperformed their life expectancy by at least 10% (RWE < 0.9), while only eight percent outperformed life expectancy to the same degree (RWE > 1.1). Importantly, this conclusion was neither inevitable nor universal. For example, in the study of *C. ibex* referenced earlier, not a single tagged animal was found to have died during their first year (Toïgo et al., 2007). In contrast, the chinook salmon (*Oncorhynchus tshawytscha*) had a first-year mortality rate of ∼94% despite a theoretical lifespan of nine years attained by a tiny proportion of individuals (Wilson, 2003). Unfortunately, this second pattern appears to be more common, and is likely to be more common in nature after taxon-related publication bias and differences in fecundity are taken into account.

It should also be noted that RWE itself merely describes the natural state of a population. It can inform population management as a descriptive statistic for prioritizing aid to particular demographics within a population, as a low RWE indicates that something about the population’s age-specific survival pattern is out of order. However, the metric should not necessarily be maximized by any possible means; for example, higher RWE could sometimes be achieved by reducing late-life welfare as opposed to increasing early-life survival. Welfare expectancy itself, which underlies RWE, should be maximized through population management. However, the average welfare expectancy of individuals may need to be traded off against the size of a population, as increasing density has potential to reduce both survivorship and welfare.

### 4.4 Welfare elasticity

A corollary of thinking about lifetime welfare in terms of expected value is the possibility of ‘bottleneck’ ages: ages where survival rate abruptly falls, which are preceded by high survivorship and followed by positive welfare expectancy. This concept is analogous to demographic elasticity, which is analyzed to identify which life stages and vital rates exercise the most control over a population’s marginal net reproductive rate (Benton and Grant, 1999). Whereas age-specific demographic elasticity depends on the parallel dynamics of survival and fecundity, welfare elasticity depends on an age’s relation to patterns of survivorship and welfare. In general, the value of increasing survival rate at a particular age depends on the proportion of individuals in the cohort surviving to reach that age and their expectation of future welfare.

Bottlenecks occurring relatively early in life, when a respectable proportion of individuals remain alive, may be promising objects for wildlife interventions from both a conservation perspective accounting for both biodiversity and welfare (Carslake et al., 2009). However, because of how few individuals of most species survive to adulthood, the conditions for a mid-life bottleneck period to be the most sensitive target for intervention appear to be uncommon. Thus, conservation interventions justified on holistic welfare grounds are likely to be most efficient when they target younger animals, who will generally be more numerous. Calculations of the expected value of any welfare intervention should account for the ages of individuals who would be affected by the intervention.

A more precise understanding of these survival and welfare parameters could elaborate on welfare expectancy through related statistical concepts, such as welfare skewness and variability (c.f. Caswell 2009 for life expectancy). Variance in welfare would be particularly important to understand if we prioritize solving cases of extremely poor welfare. If intraspecific variation in welfare is structured by geography, phenology or phylogeny, it might also be appropriate to study and manage the welfare of those groups separately, similar to how demographically independent units are often managed separately for biodiversity conservation (e.g. Höglund et al., 2011).

### 4.5 Death as a discrete welfare event

Previous publications have reasoned that for an individual animal to have had a ‘life worth living’, they must have experienced enough pleasure during their life to compensate for a potentially painful death (e.g. FAWC, 2009; Scherer et al., 2018). For animals who are able to live out most of their full lifespans, this seems highly plausible; but for the vast majority of animals, who experience only a small fraction of their potential lives, far more research into the causes and their experiences of death is needed to understand the valence of their lives.

Cause of death, and therefore the duration and pain of an animal’s experience of dying, may also vary with age similarly to welfare, though probably less systematically. In a hypothetical species, juveniles might be most likely to starve while adults are most likely to be predated, with the relative probabilities of these and other mortality factors shifting over a lifetime. If future research suggests that the pain of death is a sufficiently strong factor to negate some of the positive welfare an animal might have experienced while alive, age-specific variation in the incidence of various manners of death and their severity would also be important to account for. It is already possible to assess the welfare state of an individual - and to compare individuals within a species - using physiological and behavioral indicators. Several studies have documented consistent differences in stress hormone levels associated with different causes of death, supporting the intuitive hypothesis that some involve greater suffering than others. For example, stranded whales showed dramatically higher fecal glucocorticoid (fGC) concentrations than fishing gear-entangled whales, whose fGC concentrations were in turn dramatically higher than those of whales killed quickly by a vessel strike (Rolland et al., 2017). Similarly, deer who were shot with a rifle showed lower cortisol levels than those hunted by dogs (Bradshaw and Bateson, 2000).

### 4.6 Conclusions and implications

The consideration of age structure when evaluating the overall state of welfare in a wild animal population brings several general implications and heuristics. 1) Most individuals live only a tiny proportion of their potential lifespans, so the welfare of healthy adults, who tend to be most visible, is not representative. 2) As a consequence of this, interventions to improve welfare can normally achieve greatest impact by focusing on the youngest animals. 3) Welfare and manner of death are likely to vary with age, potentially disrupting or augmenting the focus on the youngest animals. The ideal welfare scenario - within a fixed theoretical lifespan - is for as large a proportion of animals as possible to live through the most pleasant years of life and die at the age where the typical manner of death is the quickest and least painful. 4) Since only living animals experience any welfare at all, life expectancy is a crucial factor in determining the scope for positive or negative welfare. However, if welfare varies with age, the typical individual may experience higher (or lower) net welfare than their relative life expectancy would suggest.

At the individual level, welfare expectancy unites two distinct concepts: the day-to-day quality of welfare and quantity of welfare experienced over an individual’s lifetime. However, a similar quantity-quality distinction applies at the level of populations, with welfare expectancy addressing the quality side of the argument and quantity being determined by the population size. Management decisions should be based on the sum of welfare expectancy, but density dependence of age-specific survival rates will in many cases lead to a trade-off between the average and the sum of welfare expectancy in a population (assuming habitats do not grow), implying the existence of an optimum density (e.g. Cubaynes et al., 2014). Understanding the relative sensitivities of a specific population’s vital rates to density is therefore crucial for optimal welfare-centric management.

Once better data on age-specific welfare become available, the welfare expectancy framework could also help wildlife managers to identity specific ages or stages to target for population control where a reduction in survival rate would lead to the smallest possible change in welfare expectancy for the largest possible reduction in net reproductive rate. Such compromises could also be identified for growth-oriented population management, ideally achieving high individual welfare among a large population.

The field of welfare biology is at a very early stage, having received little dedicated work from the life sciences until recently. While progress is still limited by the lack of empirical studies of wild animal welfare, it is hoped that this theoretical work, drawing on some of the same published demographic data which are widely used for informing biodiversity conservation, will help establish a paradigm for prioritizing and interpreting future research in welfare biology.

## Acknowledgements

I thank Wild Animal Initiative for their support of my research. I also thank Matthew Allcock and Brian Tomasik for their valuable feedback.

## Appendix

**Table A1:**
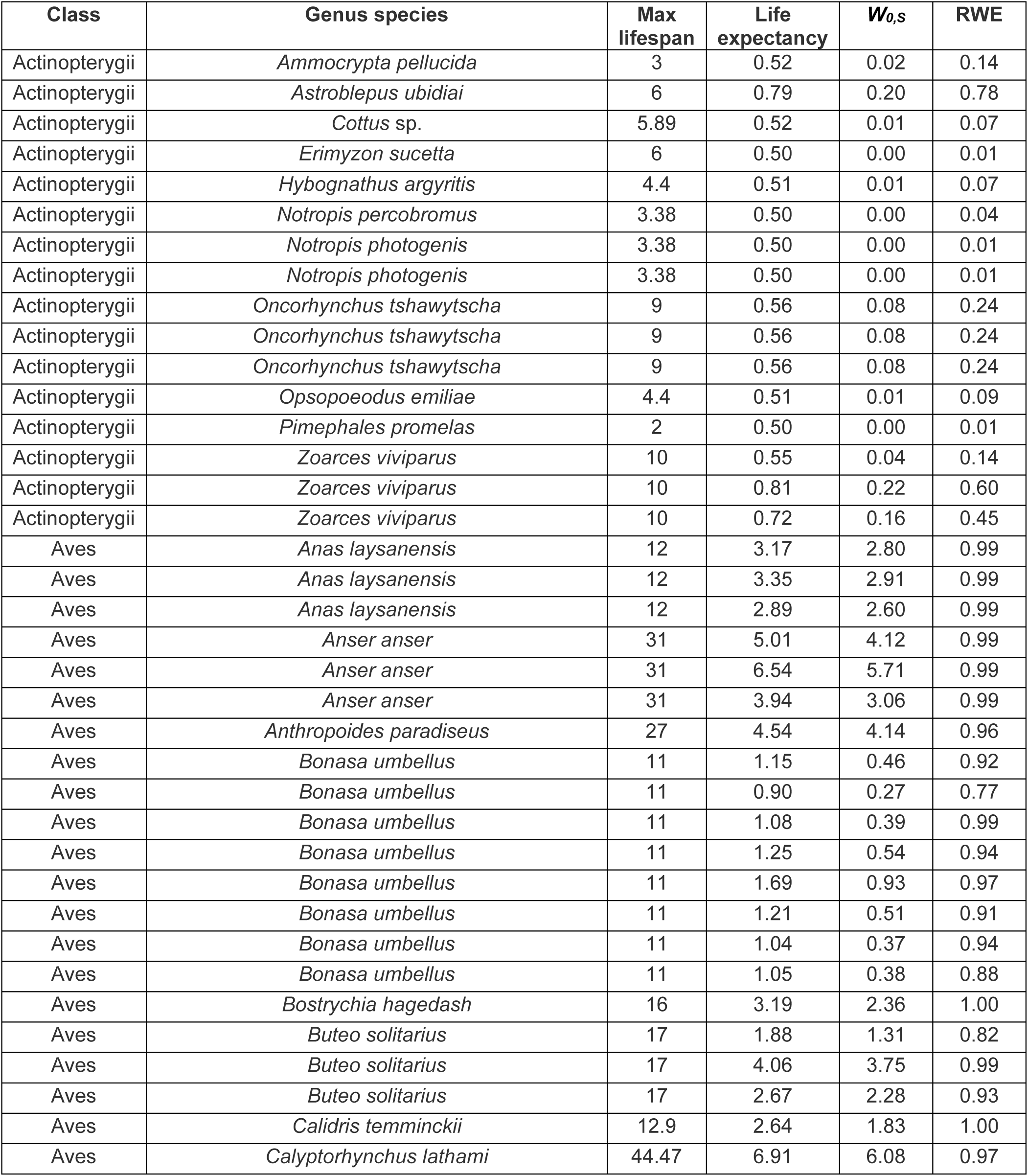
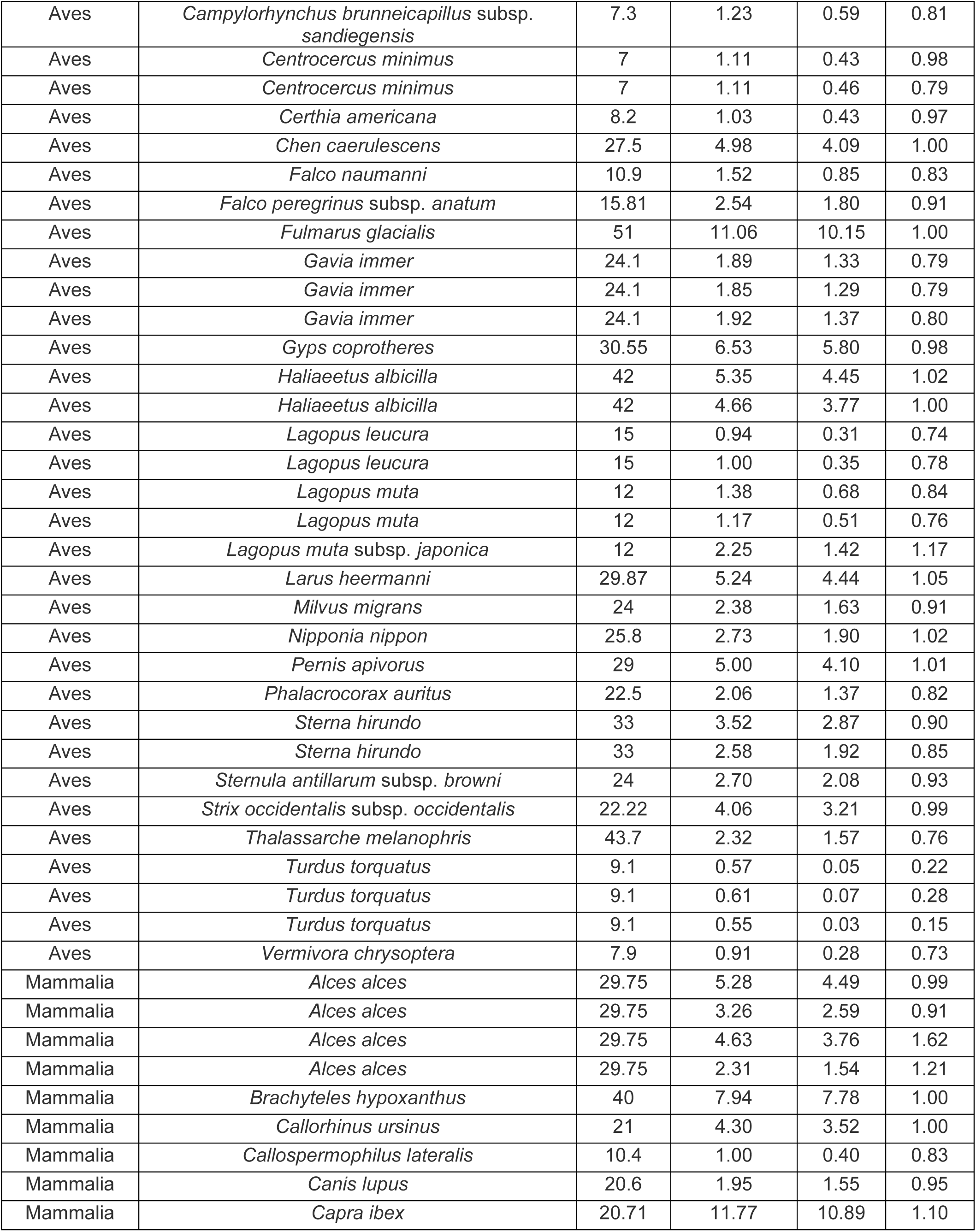
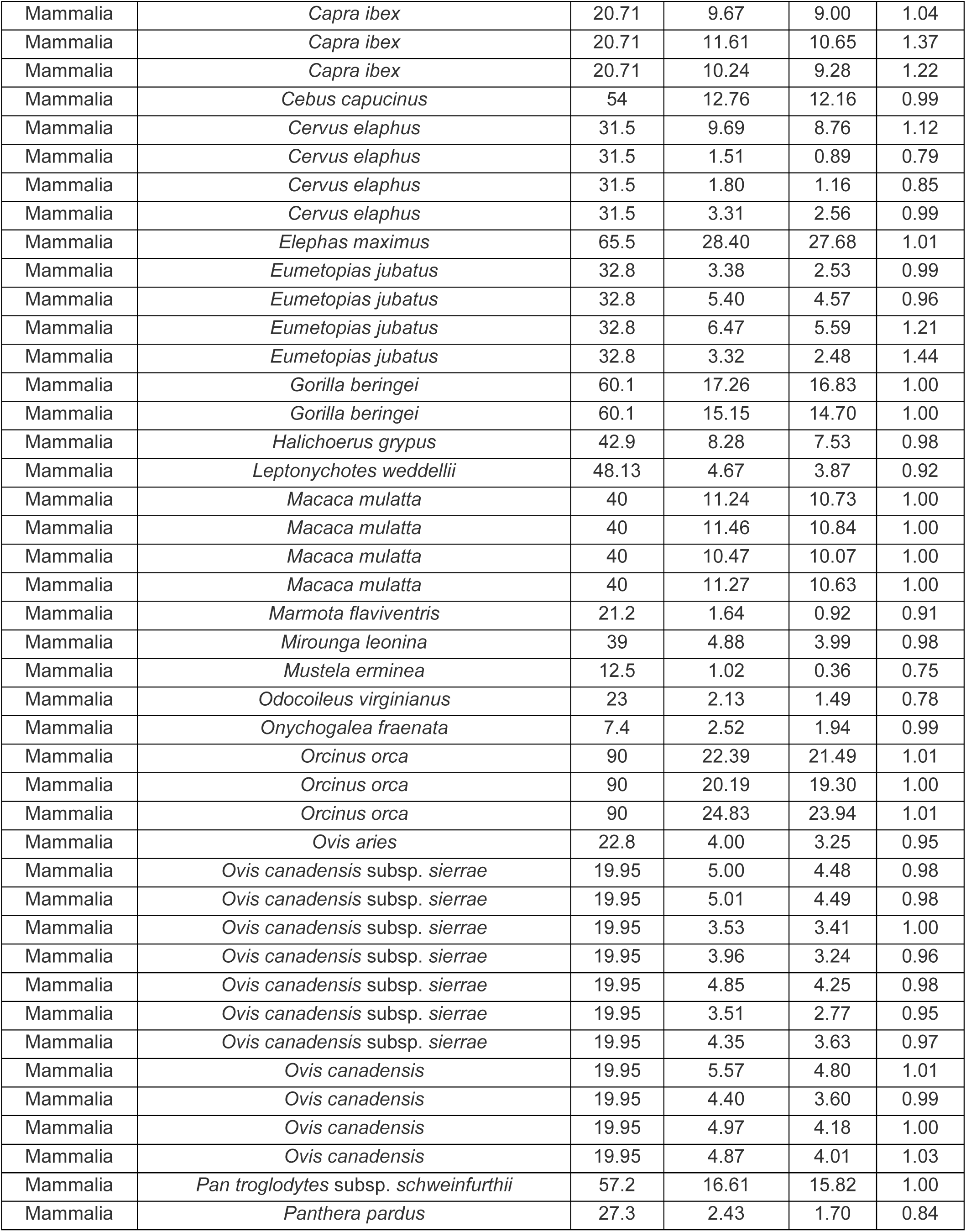
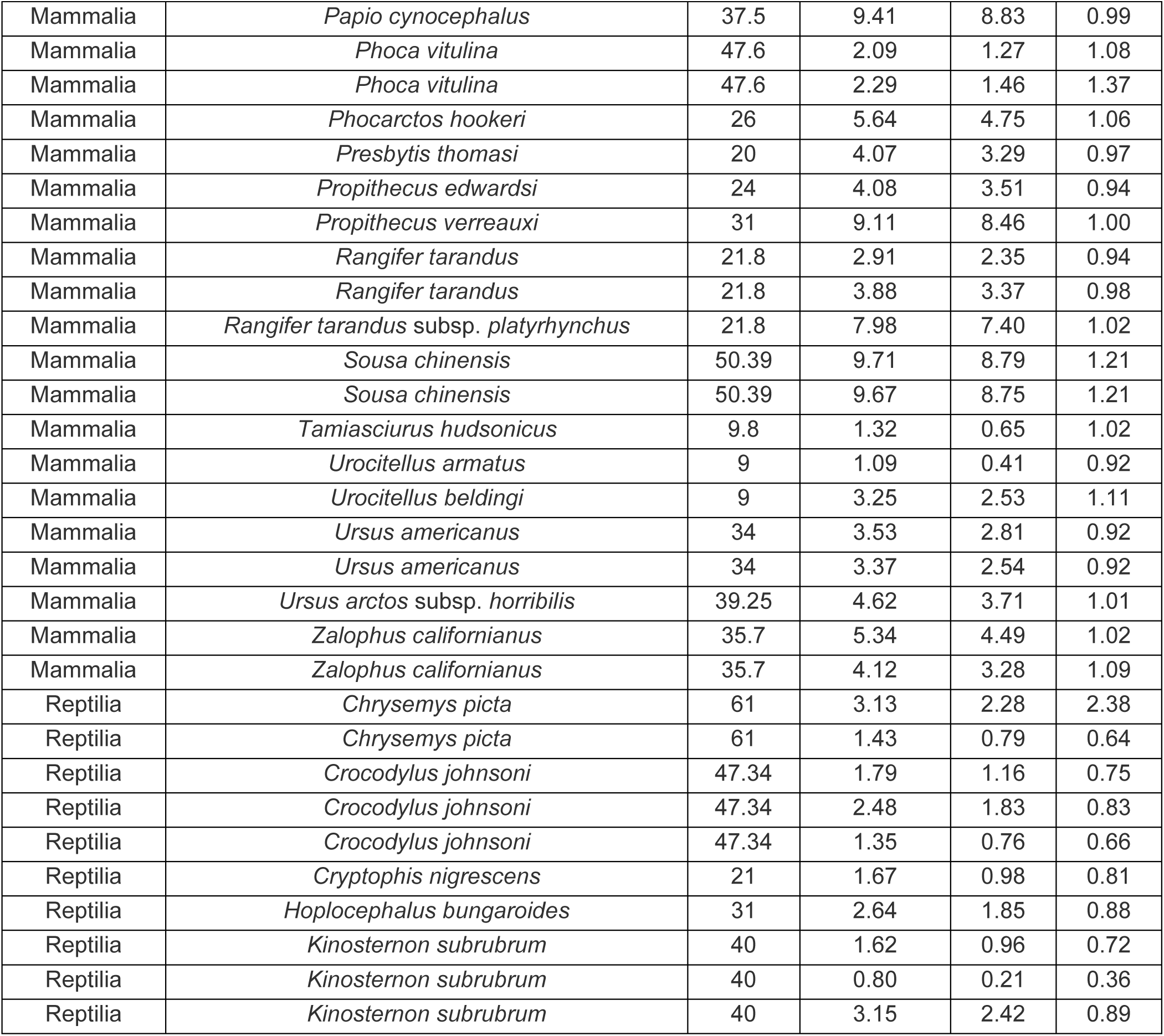
Core numerical results for each population from COMADRE included in the analysis.

